# Semi-quantitative detection of pseudouridine modifications and type I/II hypermodifications in human mRNAs using direct and long-read sequencing

**DOI:** 10.1101/2021.11.03.467190

**Authors:** Sepideh Tavakoli, Mohammad Nabizadehmashhadtoroghi, Amr Makhamreh, Howard Gamper, Caroline A. McCormick, Neda K. Rezapour, Ya-Ming Hou, Meni Wanunu, Sara H. Rouhanifard

## Abstract

We developed and applied a semi-quantitative method for high-confidence identification of pseudouridylated sites on mammalian mRNAs via direct long-read nanopore sequencing. A comparative analysis of a modification-free transcriptome reveals that the depth of coverage and specific k-mer sequences are critical parameters for accurate basecalling. By adjusting these parameters for high-confidence U-to-C basecalling errors, we identified many known sites of pseudouridylation and uncovered new uridine-modified sites, many of which fall in k-mers that are known targets of pseudouridine synthases. Identified sites were validated using 1,000-mer synthetic RNA controls bearing a single pseudouridine in the center position which demonstrate systematical under-calling using our approach. We identify mRNAs with up to 7 unique modification sites. Our pipeline allows direct detection of low-, medium-, and high-occupancy pseudouridine modifications on native RNA molecules from nanopore sequencing data as well as multiple modifications on the same strand.

## Introduction

Enzyme-mediated RNA chemical modifications have been extensively studied on noncoding RNAs^1,2^; however, messenger RNAs are also targets of RNA modification. Although modifications are less frequent in mRNAs than other RNAs^3^, these modifications potentially impact gene expression^4^, RNA tertiary structure formation^5^, or the recruitment of RNA-binding proteins^6^. Pseudouridine (ψ) is synthesized from uridine by one of more than a dozen pseudouridine synthases identified to date^7^. It was the first discovered RNA modification^8^ and represents 0.2-0.6% of total uridines in mammalian mRNAs^3^. Other uridine modifications on mRNAs include 5-methyl uridine^9^ and dihydrouridine^10,11^,but these occur to a lesser extent on mRNAs. Ψ-modified mRNAs are more resistant to RNase-mediated degradation,^12^ and have the potential to modulate splicing,^13^ immunogenicity,^14^ and translation^15,16^ *in vivo*. Further, ψ modifications of RNAs are responsive to cellular stress, leading to increased RNA half-life^17,18^. A critical barrier to understanding the precise biological functions of pseudouridylation is the absence of high-confidence methods to map ψ-sites in mRNAs. Ψ modifications do not affect Watson-Crick base pairing^19^, thereby making them indistinguishable from uridine in hybridization-based methods. Additionally, this modification bears the same molecular weight as the canonical uridine, making digested nucleotides challenging to detect directly by mass spectrometry^20,21^, and even more difficult to detect within a specific mRNA sequence from cells due to requirements of high input and purity.

Ψ is conventionally labeled using N-cyclohexyl-N’-b-(4-methylmorpholinium) ethylcarbodiimide (CMC), a reagent that modifies the N1 and N3 positions of ψ, N1 of guanine, and the N3 of uridine^22^. Treatment with a strong base removes the CMC from all the sites except for the N3 position of ψ. Recently, the use of an RNA bisulfite reaction was demonstrated for ψ-specific labeling^23,24^. Chemical labeling of ψ combined with next-generation sequencing^3,18,23^ has yielded over 2,000 putative ψ sites within mammalian mRNAs, but different methods identified different sites with limited overlap^25^, pointing to a need for alternative detection methods. Finally, since these methods rely on short reads, it is difficult to perform combinatorial analysis of multiple modifications on one transcript. Here we aim to develop a direct and orthogonal method for pseudouridine detection on mRNAs without relying on intermediate chemical reactions, but with the ability to detect previously annotated sites and to uncover new sites.

Recently, several studies have reported using nanopore-based direct RNA sequencing to directly read RNA modifications^26–31^. In these reports, ion current differences for different k-mer sequences (k = 5 nucleotides) as an RNA strand is moved through the pore suggest the presence of a modified RNA base. Detection of ψ using nanopores was also confirmed for ribosomal RNAs (rRNAs)^28^, for the *Saccharomyces cerevisiae* transcriptome^29^, and for viral RNAs^31^, as indicated by a U-to-C base-calling error at various sequence sites. Algorithms for ψ quantification have been produced^29,30^ using combinatorial sequences that contain many ψ sites within close proximity, and control RNAs that contain many natural RNA modifications also in close proximity (e.g., rRNA). While the control molecules in these studies allow analysis of many k-mers, the accuracy of quantifying ψ occupancy at a given site can be highly dependent on the nucleotide sequence surrounding the modification. Moreover, sequence context is particularly important for quantification of RNA molecules wherein the secondary structure can influence the kinetics of translocation through the nanopore^32^. Control molecules for ψ modification that match the transcriptome sequence containing the k-mer of interest are more desirable than random sequences.

Here, we describe a nanopore-based method to identify known ψ modifications and map uridine modifications in a HeLa transcriptome. We compare the sequence alignment to identical negative controls without RNA modifications and show that the number of reads and specific k-mer sequences are critical parameters for defining ψ sites and for assigning significance values based on these parameters. Our approach recapitulates 198 previously annotated ψ sites, 34 of which are detected by 3 independent methods, thus providing a “ground truth” list of ψ modifications in HeLa cells. Our approach also reveals 1,691 putative sites of uridine-modification that have not been reported previously. We show that these new sites tend to occur within k-mer sequences that are often recognized by pseudouridine synthases, such as PUS7 and TRUB1.

Applying our algorithm for detecting ψ modifications using rRNAs which have been comprehensively annotated by mass spectrometry, we assigned 41/46 ψ modifications. We demonstrate, however, that rRNA is not suitable for benchmarking RNA modifications by nanopore sequencing due to the frequent clustering of RNA modifications nearby the ψ-site, which interferes with the accuracy of basecalling, leading to false positives. Additionally, we synthesized and analyzed five 1,000-mer synthetic RNA controls containing either uridine or ψ within the sequence context of a known pseudouridylated position in the human transcriptome. This analysis revealed that U-to-C mismatch errors are systematically under-called for the detection of ψ, enabling us to identify 40 high-occupancy ψ sites, which we denote as hypermodified type I.

Further, we identify 38 mRNAs with up to 7 high-confidence ψ sites, which are confirmed by single-read analysis. Combined, this work reports a pipeline that enables direct identification and semi-quantification of the ψ modification on native mRNA molecules. The long nanopore reads allow for the detection of multiple modifications on one transcript, which can shed light on cooperative effects on mRNA modifications as a mechanism to modulate gene expression.

## Results

### Nanopore analysis of an unmodified HeLa transcriptome generated by *in vitro* transcription

To identify putative sites of mRNA ψ modifications, we extracted RNA from HeLa cells and prepared two libraries for direct RNA sequencing (**Fig. 1a**). The first (direct) library consists of native mRNAs, and the second is an *in vitro* transcribed (IVT) mRNA control library in which polyadenylated RNA samples were reverse transcribed to cDNAs, then *in vitro* transcribed back into RNA using canonical nucleotides to delete the RNA modifications. Each library was prepared for sequencing using Oxford Nanopore’s Direct RNA Sequencing Kit, and then sequenced on a separate MinION flowcell and basecalled using *Guppy 3.2.10*. Three replicates of the native RNA library produced an average of ~1.2 million poly(A) RNA strand reads, of which ~800,000 have a read quality of 7, with an average of N50 read length (defined as the shortest read length needed to cover 50% of the sequenced nucleotides) of 850 bases and a median length of ~670 bases (**Supplementary Fig. 1**). Further, we compared the coverage for individual genes in the direct RNA libraries and found that transcripts per million (TPMs) were very similar (R^2^= 1.06 for replicates 1 and 2 and 0.97 for replicates 2 and 3; **Supplementary Fig. 1**) Similarly, two replicates of the IVT library produced an average of 1.6 million passed the quality filter, with N50 of 890 and a median length of 710 bases. Alignment was performed using minimap2.17^33^ and the reads for each library were subsequently aligned to the GRCh38 human genome reference.

**Figure 1:**
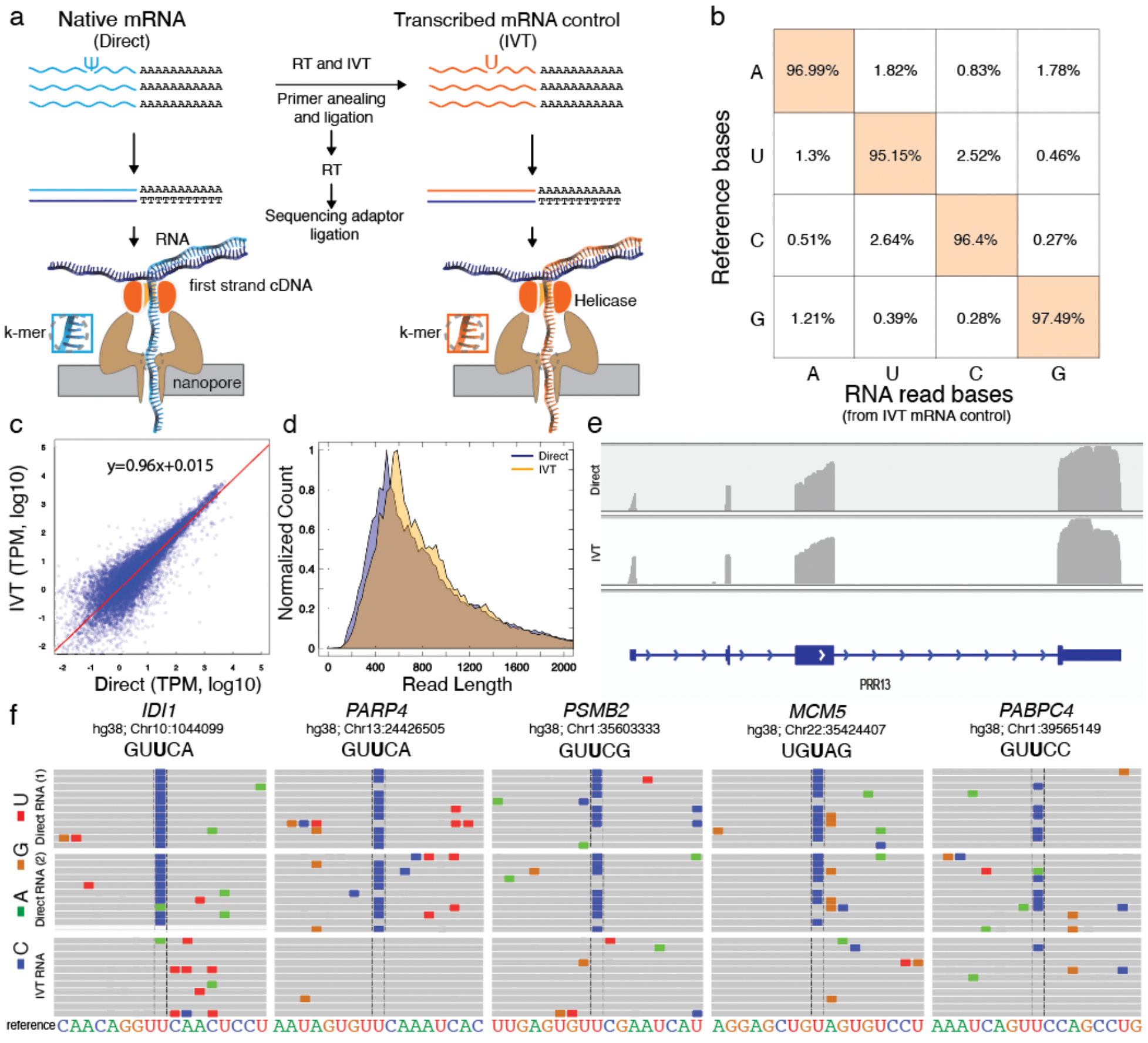
Nanopore native poly(A) RNA sequencing pipeline to identify ψ-modified sites. a. Library preparation for Nanopore sequencing of native poly(A)-containing mRNAs (direct) and sequencing of in vitro transcribed (IVT) control. b. The accuracy of called bases of *in vitro* transcribed (IVT) control samples. The x-axis shows called bases from nanopore reads and the y axis is the base identity from the reference sequence at the same position that the nanopore reads are aligned to. c. log_10_(TPM) of direct vs the log_10_(TPM) of IVT. d. Normalized count of different read lengths for direct reads (blue) vs IVT reads (orange). e. IGV snapshot of *PRR13* in direct (top) and IVT (bottom). f. Representative snapshot from the integrated genome viewer (IGV) of aligned nanopore reads to the hg38 genome (GRCh38.p10) at previously annotated ψ sites. Miscalled bases are shown in colors. Genomic reference sequence is converted to sense strand and shown as RNA for clarity.

### Basecalling accuracy is used to identify uridine modifications in RNA

To define differences between the IVT and direct libraries for uridine modification detection, any source of error other than the uridine modification itself must be minimized, including misalignments to the GRCh38 human genome reference. We minimized incorrect alignment by only considering the primary alignment of each read (i.e., the alignment with the highest mapping quality). Also, any read with a mapping quality score lower than 20 was discarded from the downstream analysis, because the probability of correct alignment was <99%. The second potential source of error is the presence of single-nucleotide polymorphisms (SNPs), whereby the base is different from the reference genome. We identified likely SNP sites based on an equivalent U-to-C mismatch percentage in both the IVT and the direct RNA sequencing samples (**Supplementary Fig. 2**). In the case of a modified RNA nucleotide, the U-to-C mismatch percentage would be significantly higher in the direct RNA sequencing sample relative to the IVT control at the site of modification (**Supplementary Fig. 2**). The third source of error is a systematic basecalling error, whereby the basecalling algorithm fails to identify the correct base. To assess the basecalling accuracy using the *Guppy 3.2.10* algorithm, we calculated the error in the IVT control by comparing the basecalling to the reference genome (**Fig. 1b**). Since the IVT control contains only the canonical RNA nucleotides, these errors were independent of RNA modifications. We confirmed that the basecaller could reliably identify unmodified and aligned nucleotides with an average U-to-C error of 2.64%.

To confirm the quality of the IVT unmodified transcriptome, we compared the coverage for individual genes in the IVT and direct RNA libraries and found that transcripts per million (TPMs) were very similar (R^2^=0.96; **Figure 1c**). We also compared the distribution of read lengths for the IVT and direct RNA libraries and found that the samples were overlapping (**Figure 1d**). Likewise, the coverage for individual transcripts was similar for IVT and direct RNA libraries (**Figure 1e**), thus validating the IVT library as an unmodified transcriptome control.

### Direct RNA nanopore sequencing identifies pseudouridines in mRNA via systematic U-to-C base-calling errors

We then examined specific locations on human mRNAs that have been previously identified as ψ sites by chemical-based methods (**Figure 1f**). We selected 5 genes as examples: *IDI1* (chr10:1044099)^3,17,23^, *PARP4* (chr13:24426505)^3,23^, *PSMB2* (chr1:35603333)^3,17,23^, *MCM5* (chr22:35424407) ^3,17^, and *PABPC4* (chr1:39565149)^3,17^, representing a range of different k-mers with a putative ψ in the center nucleotide (GU<u>U</u>CA, GU<u>U</u>CA, GU<u>U</u>CG, UG<u>U</u>AG, and GUUCC respectively). We chose a range of k-mers because specific sequences can influence the accuracy of base-calling (**Supplementary Fig. 3**). We detected a systematic U-to-C mismatch error at the reported ψ site in duplicates of each gene by direct RNA sequencing (*IDI1* (chr10:1044099):· 96.06±1.16*%*, *PARP4* (chr13:24426505): 91.71 ±7.56%, *PSMB2* (chr1:35603333): 81.07 ±1.68%, *MCM5* (chr22:35424407): 54.82 ± 4.96%, *PABPC4* (chr1:39565149): 55.08 ±3.97%). We confirmed that the IVT samples maintained the standard base-calling error at each site (3.75%, 4.54%, 1.67%, 5.26% and 8.34% respectively; **Fig. 1c**).

### Calculating the significance of U-to-C mismatch as a proxy for modification is dependent on mismatch percentage at a given site, the number of reads, and the surrounding nucleotides

To further improve the use of the U-to-C mismatch error as a proxy for U modifications we needed to minimize the error that occurs from other factors. We observed that the base quality on sites that have 3 or fewer reads is low, relative to the rest of the population, which would create bias in the downstream analysis (**Fig. 2a**). One reason for the lower quality of these sites is their proximity to the start/end of the aligned section of their corresponding reads. It is common for the aligner to clip a few mismatched bases from the start/end of reads (known as “soft-clipping”). Therefore, to ensure sufficient coverage in both the direct RNA and IVT samples, we set a minimum threshold of 7 reads represented from each biological replicate before evaluation for site modification. We show that up to 3 bases adjacent to the soft clipped site usually yield lower base quality, and thus are not reliable regions to obtain information from (**Supplementary Fig. 4**).

**Figure 2:**
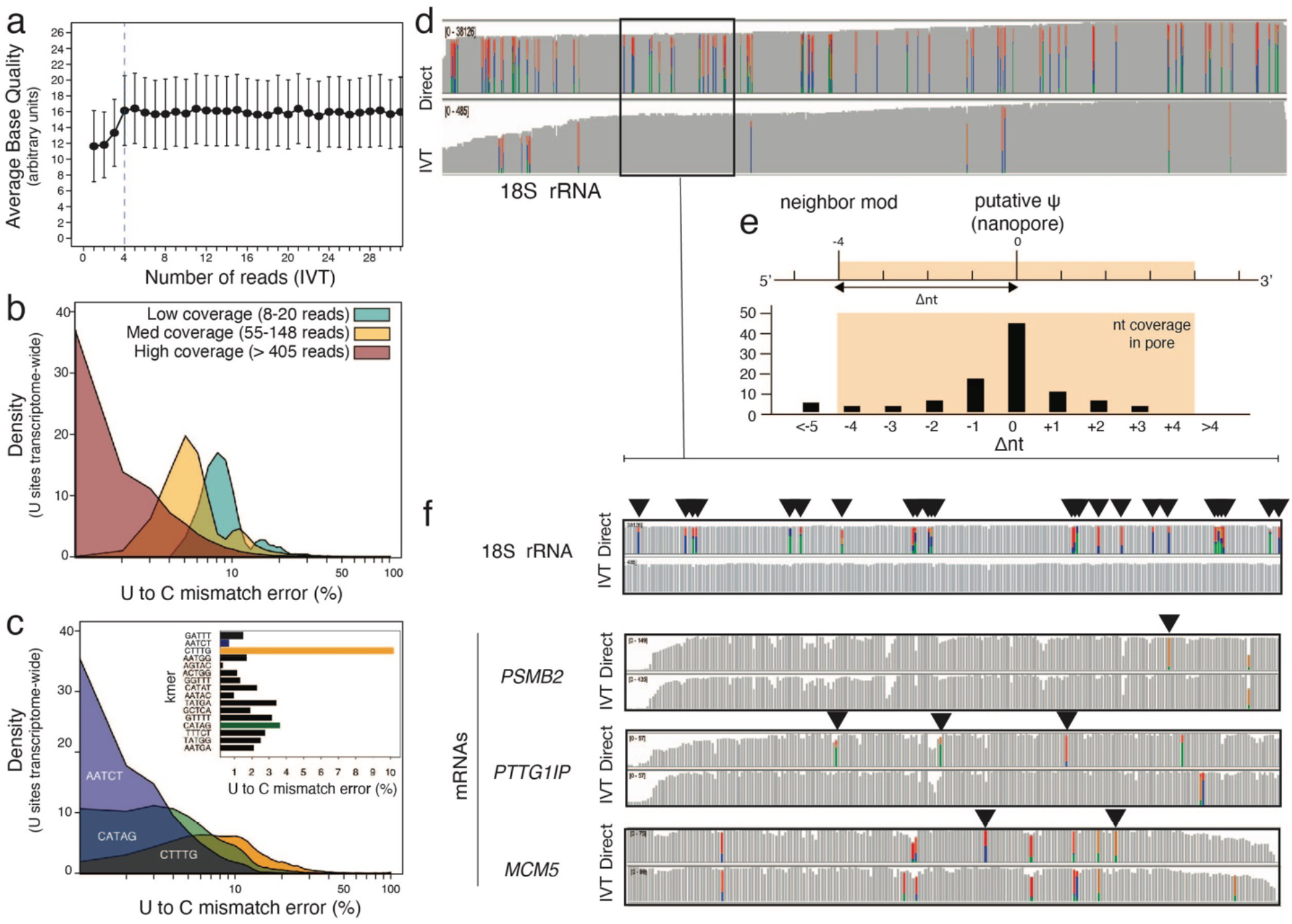
Basecalling errors can be used to detect RNA modifications if specific k-mer and coverage are considered, and the density of satellite modifications on rRNAs is different from mRNAs. a, average base quality for different numbers of reads using IVT reads. b, Distribution of U-to-C mismatch percentage for three populations, based on the read coverage. c, Distribution of U-to-C mismatch percentage for three populations, based on 5-mers. d, IGV snapshot of 18S rRNA for Direct (Upper) and IVT (lower) e, The schematic figure in which delta nt is the distance to the putative modification position. f, (top) the IGV callout of a representative 200mer section of 18S rRNA and IGV snapshots of 200mer regions within 3 mRNAs with putative modifications. Presence of a basecalling error in the Direct and not in IVT denotes a putative site of modification (**▼**).

To further investigate the U-to-C mismatch errors near the start/end of the alignment, we gathered the data for all the canonical uridine sites from our IVT control sample (>3 million uridine sites transcriptome-wide). For each of these positions, we calculated the U-to-C mismatch percentage, the number of aligned reads, and analyzed the surrounding bases of each site. We tabulated their 5-mers for which the target uridine site falls in the center. As expected, higher error rates were observed among low coverage sites (**Fig. 2b**). Additionally, the surrounding bases of a site influenced the mismatch error (**Fig. 2c**). For example, uridine sites within the CUUUG k-mer, on average, showed a 10% mismatch error in the IVT reads, while uridine sites within the AAUCU k-mer had less than 0.4% average mismatch error. The average U-to-C mismatch of the specific k-mer in IVT is an important factor to be considered because it is essential to prevent a misinterpretation of the inherent error of a k-mer as a site of modification. Therefore, the significance of the U-to-C mismatch percentage of a site must be interpreted based on a combination of the mismatch percentage (in the direct RNA sample), the number of reads (in the direct RNA sample), and the average U-to-C mismatch error of the equivalent k-mer in the IVT sample **(Methods** and **Supplementary Methods**).

It is important to ensure that the targets are not selected based on errors from other sources such as single-nucleotide polymorphisms, basecalling, or alignment. In the cases that the IVT error at a specific position is higher than the average error of that k-mer, the mismatch error from the direct RNA reads is compared with error at the specific site rather than with the average error of that k-mer in IVT. To account for standard basecalling errors, we compare the direct reads to the IVT replicate, using the highest U-to-C error at that site in the direct reads.

### Detection of annotated ψ sites (*p* < 0.001) on human rRNA

Human rRNA has been extensively annotated using mass spectroscopy and is commonly used to benchmark the specificity of RNA modification. We generated and analyzed direct rRNA and IVT rRNA libraries from HeLa cells. A total of 43 previously annotated rRNA positions from the 18S and 5.8S subunits had sufficient coverage for analysis, while the 28S in the large subunit was omitted from analysis due to insufficient coverage. Of these sites, 38 (88.4%) were detected as ψ (p < 0.001; **Figure 2d, e**, **Supplementary Table 1**). In addition to the known ψ sites that were determined by our algorithm, we detected 72 targets that are not on the list of previously detected ψ positions. Further inspection reveals that 10/72 of those positions exist in annotated rRNA positions not as ψ, but as modified 5-methyl uridine. A majority of the remaining positions (53/62) are within 4 bases of another previously annotated modification in each rRNA (**Figure 2d, e**, **Supplementary Figure 5**, **Supplementary Table 1**).

### Comparison of mismatch error as a proxy for modification density on rRNA and mRNAs

Due to the extensive annotation and quantification of ψ sites on rRNA using mass spectrometry, they are considered the gold standard in the field for RNA modification detection and quantification. However, the distance between known modifications falls within the 12-nucleotide window of the biological nanopore, thus likely disrupting expected signal patterns, as shown in **Figure 2e**. A comparison of basecalling error at sites in the direct reads and no error in the IVT control demonstrates the high density of modifications across the 18S rRNA sequence with 25 errors within a ~200 nt window (**Figure 2d, f**). In contrast, the 200 nt regions flanking putative modification sites on mRNAs *(PSMB2, PTTG1IP* and *MCM5)* have 1-4 mismatch errors in the direct RNA samples as compared to the IVT controls.

### Identification of previously annotated ψ sites on HeLa mRNA (*p* < 0.001)

Previous studies have identified putative ψ sites on human mRNA using biochemical methods including CMC^3,17,18^ and RNA bisulfite^23^ (**Fig. 3a-d**). To evaluate the accuracy of our nanopore-based method in identifying previously annotated ψ sites, we generated a list of 334 putative ψ positions focusing on those that produced a minimum of 7 reads by nanopore sequencing. We assigned *p* values (**Figure 2d**) for each of the previously annotated ψ positions and found 232 positions with *p* < 0.01 (**Fig. 3e**). Among these positions, 198 sites were previously annotated by one other method and 34 were previously annotated by two or more methods^23^ which we define as “ground truth” due to the identification of the same site by all three methods, i.e., 2+ methods and nanopore (**Fig. 3f, Supplementary Table 1**). For sites with sufficient coverage, our algorithm for determining ψ positions from direct RNA nanopore libraries had the highest overlap with Pseudo-seq (87.8%), followed by RBS-seq (77.9%), and had lowest overlap with CeU seq (67.6%).

**Figure 3:**
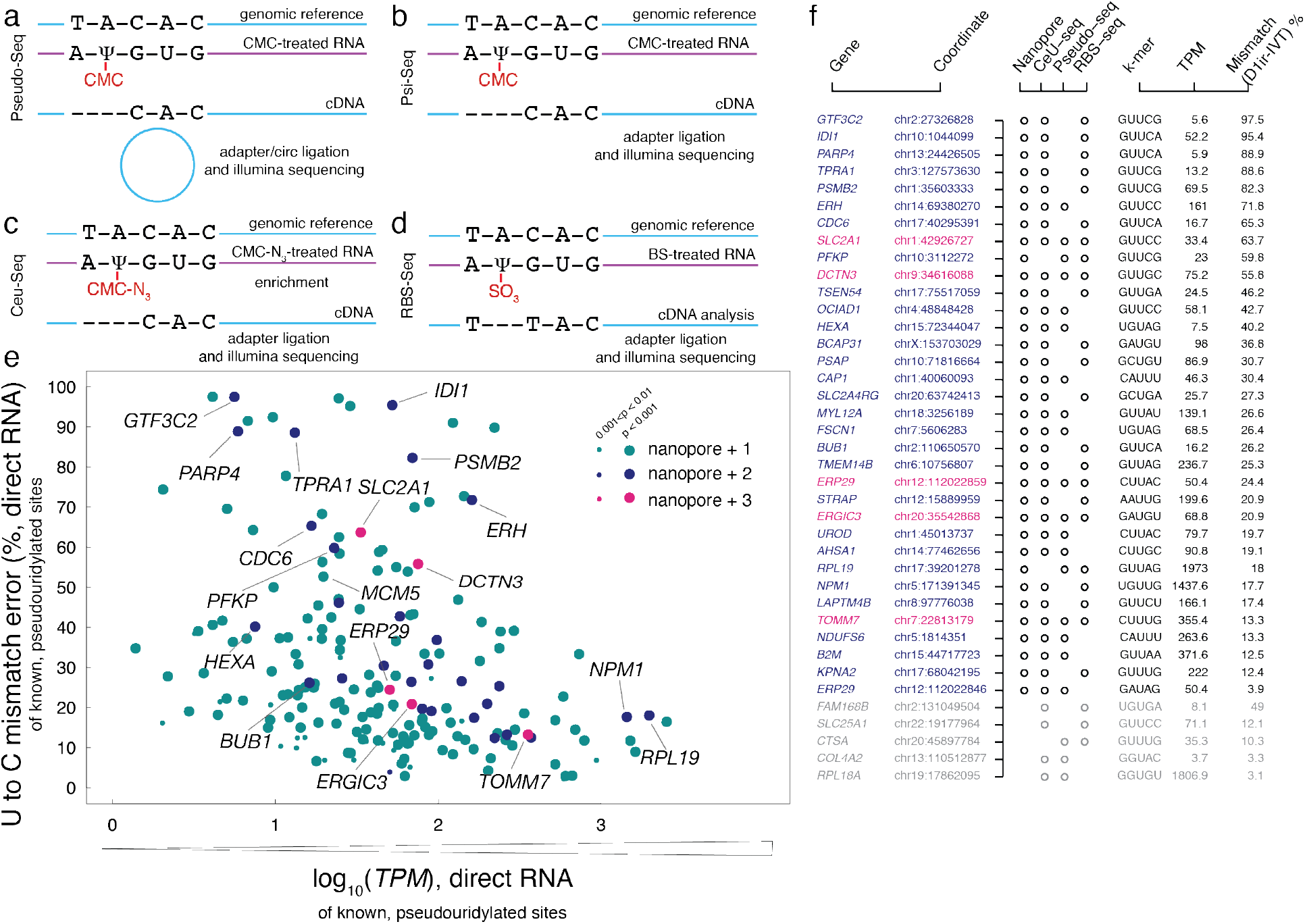
Previously annotated ψ modifications in the human transcriptome are validated by nanopore sequencing. a. The schematic workflow of the CMC-based methods that have detected ψ modification in the human transcriptome. a. Pseudo-Seq, b. Ψ-Seq, c. CeU-Seq, and d. modified bisulfite sequencing (RBS-Seq). e. U-to-C mismatch error (%) of the merged replicates of direct RNA of known ψ sites versus the log10(TPM) of merged direct RNA sequencing replicates. All targets shown are picked up by nanopore method and are previously annotated by at least one previous method. teal: annotated by one previous method, blue: annotated by two previous methods, magenta: annotated by three previous methods. f. The annotation of the genes containing a reported ψ modification by two or more previous methods. The ones validated by nanopore sequencing with a high confidence value (*p* of both replicates < 0.001) (black) and not validated by our nanopore method (Grey).

Additional analysis of the positions that were previously annotated by 2 or more independent biochemical methods revealed only 5 positions that have sufficient coverage for analysis but were not identified as having a ψ by our algorithm (**Fig. 3f**). These positions include *COL4A2* (chr13:110512877), *RPL18A* (chr19:17862095), *CTSA* (chr20:45897784), and *SLC25A1* (chr22:19177964), each of which had a low mismatch error in direct RNA sequencing. Additionally, FAM168B *(chr2:131049504)* had high error in the corresponding IVT control, indicative of k-mer specific noise within those sequence contexts.

### Transcriptome-wide detection of uridine-modified sites in human cells

Next, we sought to apply our method for *de novo* detection of transcriptome-wide uridine modifications. We broadly encompassed ψ, DHU (dihydrouridine), and possibly Um (2’-O-methyl uridine) as uridine modifications that can occur in human transcriptome^9,10^. To minimize the inclusion of sites of random error, we calculated significance based on the higher error of two replicates of analysis. We also required that two out of three direct replicates have the *p* ≤ 0.01 to be defined as a site of uridine modification. Using this algorithm, we detected 1,691 uridine-modification sites (*p* < 0.01), including 730 positions with a *p* value cutoff of 0.001 (**Fig. 4a, Supplementary Table 3**). Gene ontology analyses (GO Molecular Function 2021) were performed on genes with *p* < 0.001 using enrichR website, showing that the “RNA binding” group has the highest normalized percentage of these genes. The enrichment of “RNA binding” group is also found in the GO of all identified transcripts (**Supplementary Fig. 6**, **Supplementary Table 6**). We then determined if Um was among the identified uridine-modified sites in our nanopore method. From a meta-analysis of known Um sites previously identified in HeLa cells^9^, we found no overlap with the sites called from our algorithm, indicating that the U-to-C base-calling does not report on the Um modification.

**Figure 4:**
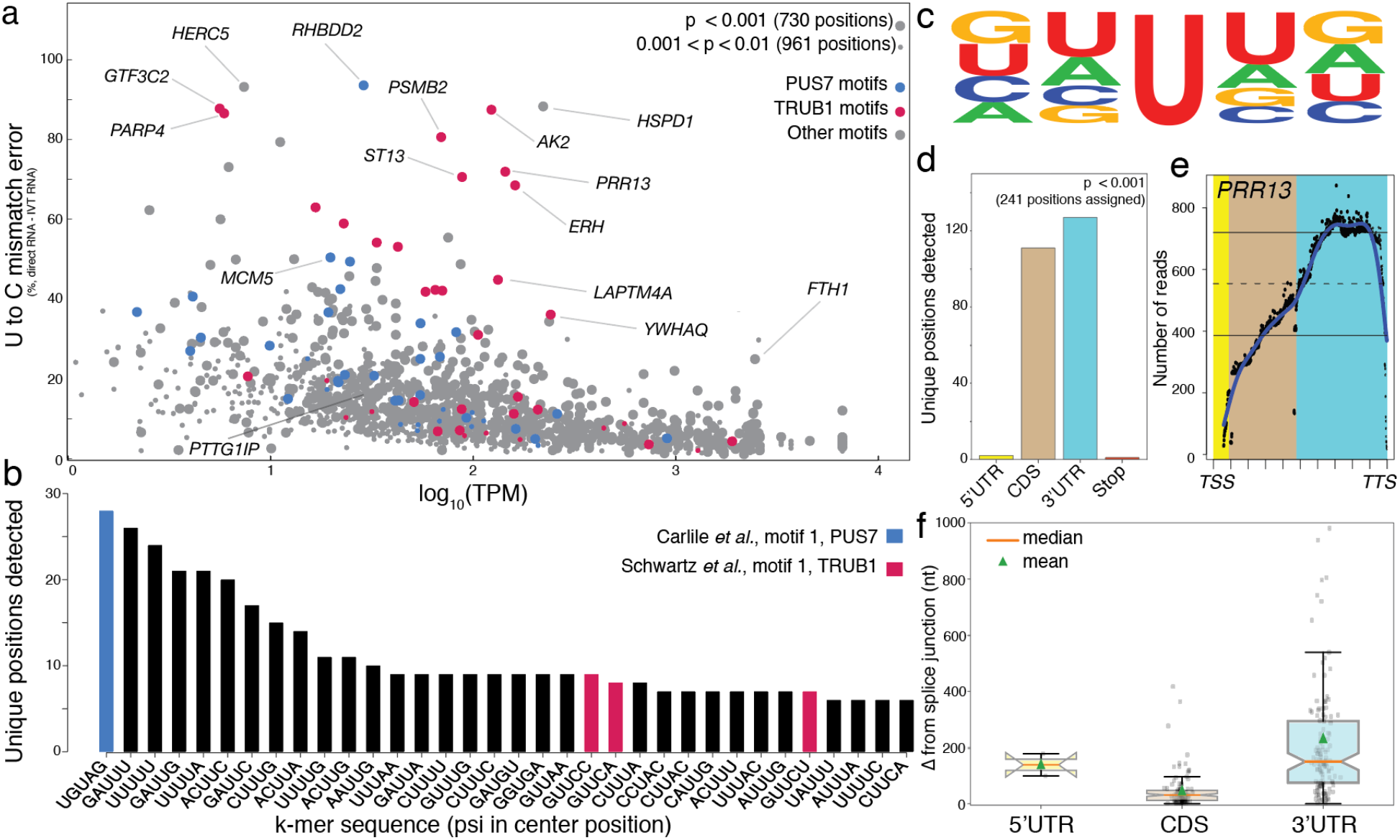
Nanopore sequencing detects uridine modifications transcriptome-wide. a. the U-to-C mismatches detected by nanopore sequencing versus the −log10(TPM) of merged direct RNA. large dot: the detected targets identified by the significance factor of two out of three replicates lower than 0.001, smaller dot: the detected targets identified by the significance factor of two out of three replicates lower than 0.01, blue: Targets with PUS7 motif, red: Targets with TRUB1 motif, and grey: Targets with the motifs other than PUS7 or TRUB1. b. The k-mer frequency of the most frequently detected targets with higher confidence. c. The sequence motif across the detected ψ modification for all detected k-mers generated with kplogo^40^. d. The distribution of detected ψ sites in the 5’ untranslated region (5’ UTR), 3’ untranslated region (3’ UTR), and coding sequence (CDS). e. The read depth of the reads aligned to PRR13 versus the relative distance to the transcription start site (TSS) and transcription termination site (TTS). f. The distance from the nearest splice junction of the sites detected in the 5’UTR, 3’UTR, or CDS after reads were assigned to a dominant isoform using FLAIR ^35^.

### Uridine-modifications are enriched within k-mer motifs of pseudouridine synthases

We assessed the k-mer frequencies for sites of uridine modification with *p* < 0.001 (**Fig. 4b**) and found that the k-mer UGUAG is the most highly represented, and that the k-mer GUUCN is among the most frequently detected. Note that UGUAG is the motif for PUS7 binding^34^, while GUUCN is the motif for TRUB1^25^, strongly indicating that they are sites of ψ modifications. To evaluate the sequence conservation of nucleotides within k-mers bearing a modification in the center position, we plotted the sequencing logo and found that the surrounding positions do not show any nucleotide preference (**Fig. 4c**).

### Distribution of putative ψ-modified sites on mRNA sequences

We characterized the distribution pattern of sites that are most likely ψ modifications on mature mRNA transcripts and observed that ~60% of them are located on the 3’ untranslated region (UTR) and 35% on the coding sequence (CDS), with very few targets detected in the 5’ UTR (**Fig. 4d**). The limited detection of ψ sites in the 5’ UTR is likely due to the low observed coverage in the 5’ end of the RNA (i.e., near the transcription start site and covering a majority of the 5’ UTR in many cases; **Fig. 4e**). Low coverage in the 5’ ends of RNA is expected since the enzyme motor releases the last ~12 nucleotides, causing them to translocate at speeds much faster than the limit of detection^26^. Compared to the rest of the transcripts, there is also a sharp drop in coverage at the tail end of the 3’ UTR, near the transcription termination site (**Fig. 4e**). Interestingly, we found one example of a putative ψ site within a transcription stop site: *GAGE2A* (chrX:49596731).

We calculated the distance of each putative ψ site from the closest splice site for high confidence ψ sites (*p* < 0.001). Prior to extracting the distance of the nearest splice junction for each target, we used the RNA isoform analysis tool, FLAIR^35^ to bin the reads comprising high confidence modified targets into their respective dominant isoform. Overall, targets in the 3’ UTRs are separated from a splice site by a longer distance relative to targets in CDS (**Figure 4f**). Considering the vast differences in sequence length between CDSs and between the 3’ UTRs, we observed a higher correlation between the splice distance of CDS-positioned targets and CDS length as compared to 3’ UTR-positioned targets (**Supplementary Figure 6**).

### Systematic under-calling of ψ percentage based on site-specific synthetic controls

To explore the quantitative potential of the U-to-C basecalling error as a proxy for pseudouridine, we constructed five synthetic mRNAs, each 1,000-mer, bearing a pseudouridine at the nanopore detected site (**Fig. 5a**). These controls were designed to recapitulate the 1,000-mer sequence flanking a natural ψ site in the human transcriptome, considering that the long-range sequence context can influence the current distributions of nanopore and basecalling for a given k-mer. Two of the chosen targets (*PSMB2;* chr1:35603333^3,17,23^ and *MCM5;* chr22:35424407^3,17^) were annotated as ψ by two or more previous methods and the other three targets *(MRPS14;* chr1:175014468, *PRPSAP1;* chr17: 76311411, and *PTTG1P;* chr21:44849705) were detected *de novo* using the U-to-C mismatch error and our *p-value* cutoff. For each site, we constructed a pair of RNA transcripts, in which the center position of the k-mer is either a uridine or a pseudouridine. We ran these synthetic controls through the nanopore directly and measured the U-to-C mismatch error for each. If the mismatch error were a perfect proxy for ψ, we expected to see 100% U-to-C mismatch in these synthetic controls. In contrast, we observed 38.17% U-to-C mismatch error for *PSMB2*, 32.16% for *MCM5*, 69.64% for *PRPSAP1*, 69.35%% for *MRPS14*, and 30.08% for *PTTG1P* (**Fig. 5b**). These results indicate a systematic under-calling of ψ using our algorithm.

**Figure 5:**
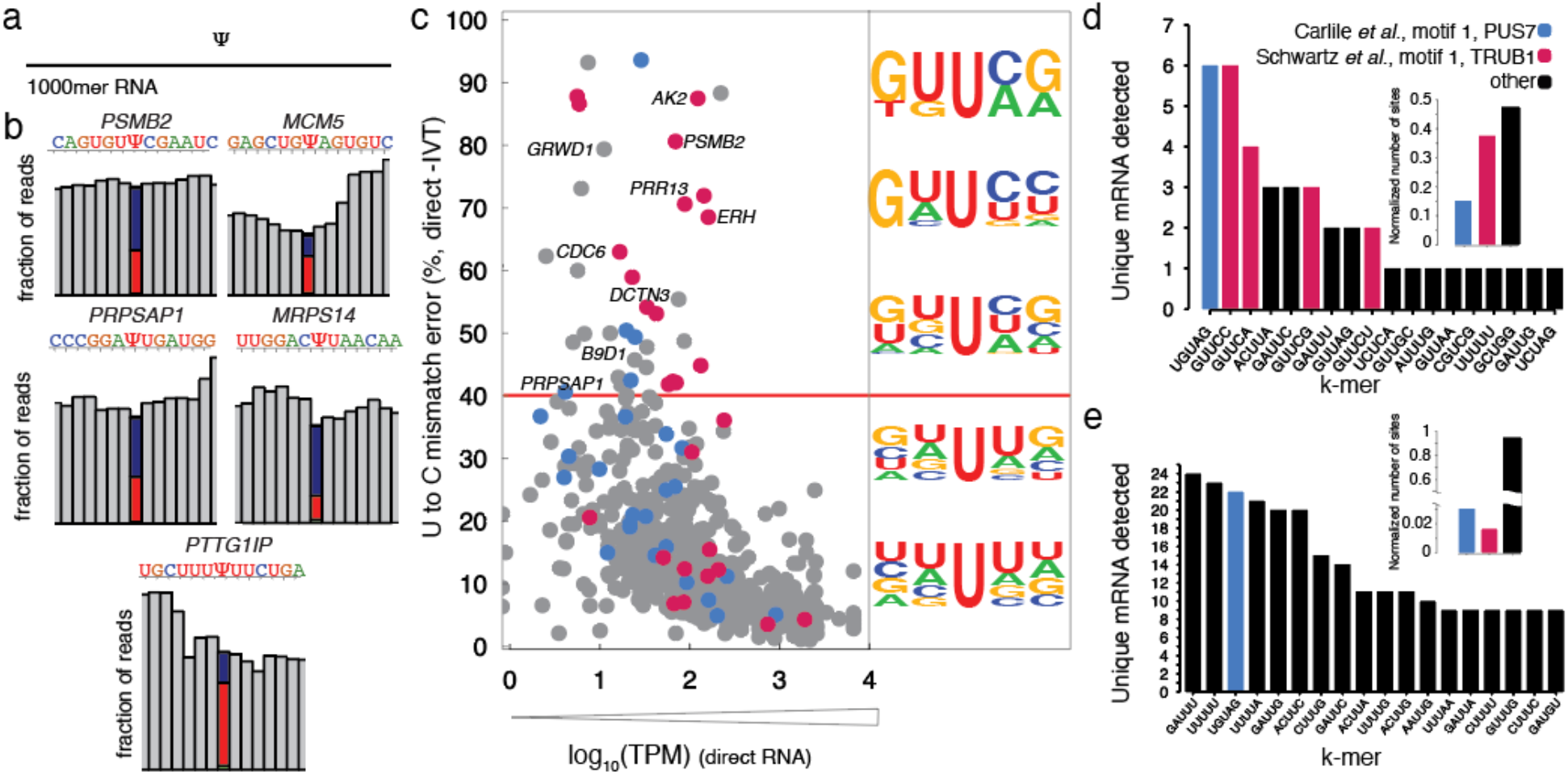
Solid-phase synthesis of 1,000-mer RNA standards that maintain the sequence context of putative ψ sites demonstrates a systematic undercalling of the modification. a. A pair of 1,000-mer synthetic RNA oligos were designed, one containing 100% uridine and the other containing 100%ψ in a sequence that recapitulates the natural occurrence of ψ in the human transcriptome. b. The frequency histograms of 13 nucleotides surrounding the detected ψ position in the middle of a k-mer in 4 different mRNAs: *PSMB2, MCM5, PRPSAP1*, and *MRPS14, and PTTG1IP*. c. The U-to-C mismatches of the detected ψ position for merged replicates of direct RNA seq versus −log10(significance). The targets with U-to-C mismatch of higher than 40% are defined as hypermodified type 1. The sequence motifs for different mismatch ranges are shown. d. K-mer frequency is shown for hypermodified type I and “not hypermodified”ψ sites with the highest occurrence. e. Distribution of U-to-C mismatches higher than 40% in mRNA regions.

### Positions with >40% U-to-C mismatch error are classified as type I hypermodification

We define hypermodification type I as a specific site within a transcript in which at least every other copy has a uridine modification. We therefore reasoned that a 40% mismatch error was an appropriate cutoff because the base caller is systematically under-calling the modification. We further reasoned that, while somewhat arbitrary, at 40% mismatch error, we encompass all of the sites with 50% occupancy and above. From our *de novo* uridine-modification detection analysis, we identified 40 unique sites of hypermodification type I including *AK2* (chr1: 33014553), *IDI1*(chr10:1044099), *GTF3C2*(chr2:196789267), *RHBDD2* (chr7:75888787)*, HSPD1 (chr2:197486726)* that show close to 100% mismatch error (**Supplementary Table 4**).

To assess the sequence conservation of nucleotides within k-mers bearing a putative ψ modification in the center position, we selected all unique sites with a U-to-C mismatch error above 40% (**Supplementary Table 4**). We found that the −1 position has a strong preference for uridine and the −2 position has a strong preference for guanosine. This preference becomes more significant as the mismatch percentage increases (**Fig. 5c**). The +1 position has a strong preference for cytosine especially at sites with higher than 80% U-to-C mismatch error.

We then assessed the k-mer frequencies for detected positions with a U-to-C mismatch error at greater than 40% (**Fig. 5d**) and those with an error less than 40% (**Fig. 5e**). Among k-mers with a frequency greater than 40% mismatch error, the GU<u>U</u>CN k-mer, representing the TRUB1 motif, is the most prominent (30/105 sites around 29%). The k-mer UG<u>U</u>AG, representing the PUS7 motif, is also prominent (5/105 sites around 4.8%). In contrast, the k-mers UG<u>U</u>AG (13/712, 1.8%), GU<u>U</u>CN, and all others occurred at a similar frequency as the most abundant “not hypermodified” targets (15/712, 2.1%).

We assessed the location of putative ψ on the transcript and found that type I hypermodified sites are biased towards 3’ UTRs, which is the same as sites that are not hypermodified (**Supplementary figure 7**). No significant difference was observed in the splice distance of type I hypermodified sites between sites in the 3’ UTR and those in CDS regions of mRNA when compared to “not hypermodified” sites (**Supplementary Figure 7**).

### Messenger RNAs with >1 uridine modifications are classified as type II hypermodification

We define hypermodification type II as the mRNAs that can be modified at two or more positions. Using only the sites with a high probability of modification (*p* <0.001), we identified 104 mRNAs with modifications at 2 unique positions, 27 with 3 positions, 4 with 4 positions, 5 with 5 positions, 1 with 6 positions and 1 mRNA with 7 positions (**Fig. 6a**). For the mRNAs that are modified at 2 positions, we found no correlation between the mismatches in position 1 and position 2 (R^2^ = 0.039), as expected because this error is highly dependent on the k-mer sequence. To demonstrate future applications of long read sequencing for assessing modifications on the same transcript, we plotted every read for two mRNAs *(CHTOP* and *PABPC4*) and labeled each site using the called base (**Fig. 6b**). We observed that these mismatches could happen on the same read. For example, 17% and 7% of the reads had a U-to-C mismatch in both positions for *CHTOP* and *PABPC4* respectively. We plotted the distribution of type II hypermodifications across the body of the transcript and found a slight clustering of sites in the 3’ UTR (**Fig. 6c**).

**Figure 6:**
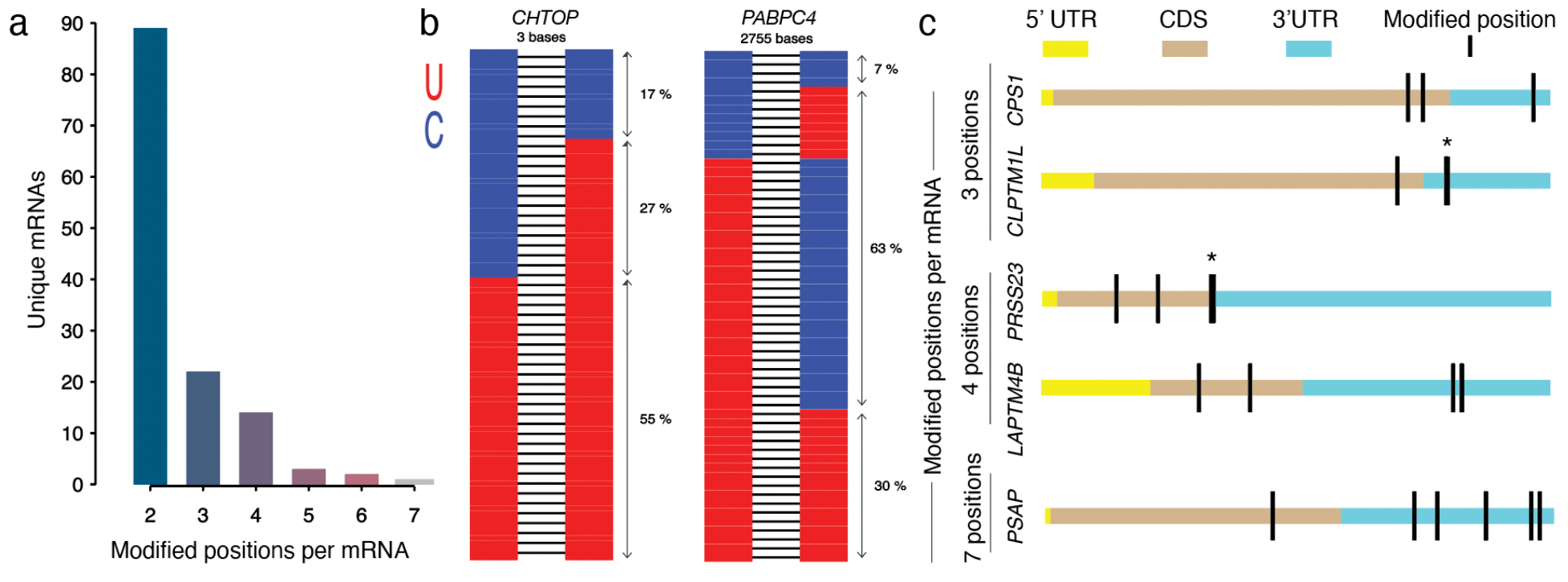
Type II hypermodification is defined as the mRNA targets that contain two or more uridine-modified positions. a. Unique mRNAs that are classified as hypermodification type II positions and the number of modified positions possible on each. b. Two examples of hypermodified type II transcripts *(CHTOP* – chr1:153,645,392-153,654,395 & *PABPC4* –chr1:39,562,394-39,565,149) with two modified positions indicating U-to-C mismatch on a single read for long reads that cover both positions. c. Examples of type II hypermodification with three or more modified positions distributed across each gene.

## Discussion

We show here that systematic U-to-C basecalling errors detected from direct nanopore sequencing of transcriptomes can serve as indicators for annotated sites of ψ modification. In this work, we provide a foundation for identifying modified U sites with high confidence based on two approaches. In the first, U-to-C mismatch errors in native transcriptomes are compared against a corresponding unmodified transcriptome as a negative control to eliminate standard basecalling errors that occur in canonical bases. We weigh the transcriptome wide average of U-to-C errors in k-mers to minimize false positives due to low coverage that is an issue with direct mRNA sequencing. In the second, we use a set of long synthetic RNA controls with precisely positioned ψ modifications to aid in our discovery of systematic under-calling of ψ modifications, pointing to limitations of basecalling-guided RNA modification detection algorithms. Our approach is distinct from the ELIGOS algorithm^36^, primarily because we consider the average U-to-C error of unmodified k-mers, enabling analysis of low coverage sites that may show as significant error due to random nanopore basecalling error. Additionally, we determine individual modification sites by the exclusive presence of U-to-C mismatches rather than including all other substitutions, deletions, and insertions at a given site; thus, reducing false positive detection.

We demonstrate that this method can identify a majority of ψ sites that were detected by CMC and bisulfite-based next-generation sequencing platforms. Importantly, we produce a “ground truth” list of ψ positions in mRNAs that are detected by nanopore and by at least one other method that has previously annotated ψ in HeLa cells. This ground-truth list consists of 198 mRNA positions detected by nanopore and one other method, and 34 positions detected by nanopore and two other methods (**Figure 3**), constituting a conservative list of targets that we suggest biologists focus on for performing functional analysis. This is a significant advance in the field because the current methods to identify ψ primarily rely on CMC modification and there’s a danger in orthogonal methods relying on the same chemical. Although alternative chemical methods have been introduced, the overlap among them is relatively low. Our demonstration that nanopore sequencing can detect modifications at the same site as chemical methods strengthens the identification at each site. The probability of a ψ site within the transcriptome being identified both by chemical methods and by nanopore sequencing is very low, because our nanopore sequencing identified only 1,691 uridine-modified sites in the transcriptome. This demonstrates the rigor of our approach and suggests that the “ground truth” list can serve as a first step for developing biological assays to determine the specific function of individual ψ modifications in mRNAs.

Among the methods that we used to validate our data, Pseudo-seq shows the highest overlap between the detection targets. However, some targets that the other methods detected were not detected by our method. We demonstrate that some of these are due to problematic k-mers that show a high error in the IVT control, or very low coverage, thereby not meeting the criteria for being called as a modified site. Alternatively, artifacts from CMC labeling may account for this, including incomplete CMC adduct removal from unmodified uridines, reverse-transcriptase read-through of CMC-modified ψ sites, or uneven amplification of low-occupancy ψ-sites. Another potential reason for the differences could be batch differences between cell lines leading to differential occupancy at a given moment of mRNA extraction. We also observed several targets that were detected by our nanopore method that were not detected by other methods. While we are confident that these sites are modified due to differences between the native RNA versus the IVT control, we cannot rule out the possibility of other uridine modifications (as shown in the identification of 5’-methyl-uridines from the rRNA data). Nonetheless, many of these targets are likely ψ due to the enrichment of k-mer sequences that are motifs for ψ synthases PUS7 and TRUB1. Further studies are necessary to resolve this issue.

However, we cannot rule out other types of uridine modifications, such as DHU^10,11^ or Um^9^ that can occur in mRNA. The estimated frequency and stoichiometry of DHU in eukaryotic mRNAs is low (<0.01%^10^) relative to ψ (0.2-0.6%^3^), but this modification also falls within uridine-rich tracks^10^. Therefore, we recommend that these uridine-modified sites be further explored as putative DHU modifications using appropriate DHU-seq^10^. While the frequency of Um is comparable to Ψ (0.15%^9^), we performed a meta-analysis of known Um sites from HeLa cells^9^ and found no overlap with the sites called from our algorithm and *de novo* analysis, indicating that the U-to-C basecalling in nanopore sequencing likely does not report on Um. Overall, our U-to-C basecalling errors represent an upper limit of ψ modification that may include other types of uridine modifications, particularly if an identified site matches with the recognition motif of one of the known ψ synthases and is confirmed by enzymatic knockdown. A better discrimination among different types of uridine modifications occurring as a U-to-C mismatch from direct RNA nanopore sequencing will require additional synthetic controls and improved machine-learning algorithms. Finally, parallel analysis of a transcriptome of interest with the transcriptome of a KO of KD cell line lacking one of the ψ synthases will help to assign a specific modification *de* novo for putative Ψ sites^17,18,29^.

Previous studies have demonstrated the importance of long-range interactions^31^ for accurate calling of ψ modifications by direct RNA sequencing. To account for the contributions of long-range interactions, we have validated our method by analysis of five synthetic 1,000-mers, each containing a site-specific ψ found within a natural target sequence in the human transcriptome. We find that the U-to-C basecalling error is systematically under-called at the modified site. Based on this finding, we defined hypermodification type I as sites that have >40% U-to-C mismatch error. We also define hypermodification type II as mRNAs bearing multiple U modification sites in a specific transcript. Finally, we show for the first time that U modifications can occur up to 7 times on a single transcript.

A fully quantitative measure of ψ occupancy at a given site would require high-coverage sequencing runs of a comprehensive set of every possible ψ-containing k-mer within its natural sequence context (an estimated 13 nucleotides surrounding the modified site). While high-throughput controls have previously been generated^29,31^, all Us were modified in those studies and consequently, these are not the ideal controls for detection of single ψ modifications within the natural sequence contexts as demonstrated in the comparison of rRNA modifications to mRNA modifications (**Figure 2**). Additionally, rRNA-based controls, which contain high frequencies of conserved modification sites, are not ideal for assessing mRNA modifications, because the spatial distribution of modifications in rRNA is much denser than that in mRNA (see **Figure 2**). Although synthetic controls are most critical to resolve ambiguities of ψ detection by nanopore sequence, the preparation of a large number of such molecules by individual synthesis of each is not feasible for a single laboratory. Future work will consider innovative methods to make libraries of all putative sites within their sequence contexts in order to quantitatively evaluate ψ profiles in transcriptomes.

Although our method is semi-quantitative, the synthetic controls that we have generated demonstrate that the basecalling error is reliable in the calling of ψ at a given site. By setting a cutoff in U-to-C mismatch for a given site we conservatively draw a list of high-confidence sites that are pseudouridylated with high occupancy, and thus, have a higher likelihood of leading to a measurable phenotype in the cell and conferring a functional impact on the cellular physiology. Our work provides a powerful foundation for detection and analysis of ψ and other uridine modifications on mRNAs with sequence specificity and single-molecule resolution. Future work should include an expansion of synthetic controls and training of a new basecaller to improve our ability to validate and quantify U modifications in transcriptomes.

## Methods

### Cell culture

HeLa cells were cultured in Dulbecco’s modified Eagle’s medium (Gibco, 10566024), supplemented with 10% Fetal Bovine Serum (FB12999102, FisherScientific) and 1% Penicillin-Streptomycin (Lonza,17602E). To extract sufficient poly-A RNA, three confluent, 10cm dishes were used for each experiment.

### Total RNA extraction and Poly(A) RNA isolation

The total RNA extraction protocol was performed using a method that is the combination of total RNA extraction using TRIzol (Invitrogen,15596026) and PureLink RNA Mini Kit (Invitrogen, 12183025). Cell types were washed with 3 ml ice-cold PBS. 2 ml of TRIzol was added to each 10cm dish and incubated at room temperature for 5 min. Every 1 ml of lysed cells in TRIzol was transferred to a LoBind Eppendorf tube and vortexed for 30 sec. 200 *μ*l chloroform (Acros Organics,423555000) was added to each tube and mixed by shaking for 15 sec and incubated at room temperature for 3 min. Then the samples were centrifuged at 12000 XG for 15 min at 4°C. 0.4 ml of aqueous supernatant is transferred to a new LoBind Eppendorf tube and an equal volume of 70% ethanol is added to the solution followed by vortexing. In the following steps, PureLink RNA Mini Kit (Invitrogen, 12183025) and the protocol are performed according to the manufacturer’s recommended protocol. Briefly, the solution is transferred to a pure link silica spin column and flow-through was discarded (every two microtubes were loaded on one column). The columns were washed with 0.7 ml of wash buffer I once and then with 0.5 ml wash buffer II twice. The total RNA was eluted using 50 ul nuclease-free water. The RNA concentration was measured using a NanoDrop 2000/2000c Spectrophotometer.

NEBNext Poly(A) mRNA Magnetic Isolation Module (E7490L) is used to select poly(A) mRNA. The protocol is followed according to the manufacturer’s protocol. The only modification was pooling 5 samples and performing the experiment in microtubes instead of PCR tubes. 15 samples (3 microtubes) were used in each experiment to get enough Poly-A RNA product. The products were eluted from the NEBNext polyA magnetic isolation (NEB, E7490S) in tris buffer. The three samples were pooled and ethanol precipitated to get to the concentration that is required for the sequencing step.

### In vitro transcription, capping, and polyadenylation

cDNA-PCR Sequencing Kit (SQK-PCS109) kit was used for reverse transcription and strand-switching. Briefly, VN primer (VNP) and Strand-Switching Primer (SSP) were added to 50 ng poly-A RNA. Maxima H Minus Reverse Transcriptase (Thermo scientific, EP0751) was used to produce cDNA. IVT_T7_Forward and reverse primers were added to the product and PCR amplified using LongAmp Taq 2X Master Mix (NEB, M0287S) with the following cycling conditions: Initial denaturation 30 secs @ 95 °C (1 cycle), Denaturation 15 secs @ 95 °C (11 cycles), Annealing 15 secs @ 62 °C (11 cycles), Extension 15 min @ 65 °C (11 cycles), Final extension 15 mins @ 65 °C (1 cycle), and Hold @ 4 °C. 1 μl of Exonuclease 1 (NEB, M0293S) was added to each PCR product and incubated at 37C for 15 min to digest any single-stranded product, followed by 15 min at 80C to inactivate the enzyme. Sera-Mag beads (9928106) were used according to the Manufacturer’s protocol to purify the product. The purified product was then in vitro transcribed using “HiScribe T7 High yield RNA Synthesis Kit (NEB, E2040S) and purified using Monarch RNA Cleanup Kit (NEB, T2040S). The product was eluted in nuclease-free water and poly-A tailed using E. coli Poly(A) Polymerase (NEB, M0276). The product was purified once again using an RNA Cleanup Kit and adjusted to 500 ng polyA RNA in 9 ul NF water to be used in the Direct RNA library preparation.

For rRNA IVT, total RNA was poly-A tailed using E. coli Poly(A) Polymerase (NEB, M0276) and purified using RNA Cleanup kit (NEB, T2040S) then poly-A selected using NEBNext polyA magnetic isolation (NEB, E7490S). 50 ng of the poly-A tailed total RNA was the in vitro transcription according to the above protocol.

### Synthetic sequence design

We constructed five synthetic 1,000-mer RNA oligos, each with a site-specifically placed k-mer. Two versions of each RNA were prepared, one with 100% uridine and the other with 100%ψ at the central position of the k-mer. The uridine-containing RNAs were prepared byT7 transcription from G-block DNAs (synthesized by Integrated DNA Technologies), whereas the ψ-containing RNAs were prepared by ligation of left and right RNA arms (each 500 nts in length) to a 15-mer RNA bearing a ψ in the central position (synthesized by GeneLink). A T7 promoter sequence with an extra three guanines was added to all the DNA products to facilitate *in vitro* transcription. In addition, a 10 nt region within 30 nt distance of ψ was replaced by a barcode sequence to allow parallel sequencing of the uridine- and ψ-containing samples. Finally, each left arm was transcribed with a 3’ HDV ribozyme that self-cleaved to generate a homogeneous 3’-end. Fulllength RNA ligation products were purified using biotinylated affinity primers that were complementary to both the left and right arms.

### Direct RNA library preparation and sequencing

The RNA library for Direct RNA sequencing (SQK-RNA002) was prepared following the ONT direct RNA sequencing protocol version DRCE_9080_v2_revH_14Aug2019. Briefly, 500 ng poly-A RNA or poly-A tailed IVT RNA was ligated to the ONT RT adaptor (RTA) using T4 DNA Ligase (NEB, M0202M). Then the product is reverse transcribed using SuperScript™ III Reverse transcriptase (Invitrogen, 18080044). The product was purified using 1.8X Agencourt RNAClean XP beads, washed with 70% ethanol and eluted in nuclease-free water. Then the RNA: DNA hybrid ligated to RNA adapter (RMX) and purified with 1X Agencourt RNAClean XP beads and washed twice with wash buffer (WSB) and finally eluted in elution buffer (ELB). The FLO-MIN106D was primed according to the manufacturer’s protocol. The eluate was mixed with an RNA running buffer (RRB) and loaded to the flow cell. MinKnow (19.12.5) was used to perform sequencing. Three replicates were from difference passages and different flow cells were used for each replicate. For Direct rRNA library preparation, total RNA was poly-A tailed using E. coli Poly(A) Polymerase (NEB, M0276) and purified using RNA Cleanup kit (NEB, T2040S) following up with the above protocol.

### Base-calling, alignment, and signal intensity extraction

Multi-fast5s were base-calling real-time by guppy (3.2.10) using the high accuracy model. Then, the reads were aligned to the genome version hg38 using minimap 2 (2.17) with the option ‘‘-ax splice -uf -k14’’. The sam file was converted to bam using samtools (2.8.13). Bam files were sorted by “samtools sort”and indexed using “samtools index”and visualized using IGV (2.8.13). The bam files were sliced using “samtools view -h -Sb” and the signal intensities were extracted using “nanopolish eventalign”.

### Gene ontology and sequencing logo analysis

Gene ontology (GO) analysis of Molecular Function 2021 was performed using enrichR website ^37–39^. The sequence motifs are generated by kpLogo website ^40^.

### Modification detection and analysis

A summary of the base calls of aligned reads to the reference sequence is obtained using the *Rsamtools* package. Mismatch frequency is then calculated for a list of verified ψ sites. We observe that U-to-C mismatch frequency shows a better separation between the modified (IVT) and (potentially) modified (Direct) samples (refer to the scatter plots from SI, talk about the p-value from t-test that will be included for each panel in the caption).

We know from our control sample that U-to-C mismatch frequency depends on both the molecular sequence and coverage (**Fig 2. a, b, and c**). Therefore, the significance of an observed mismatch percentage at each site is calculated accordingly and via the following equation:

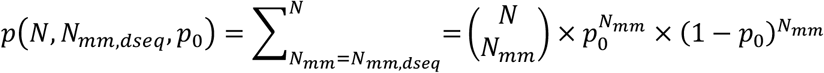

where the significance of the mismatch frequency at each U site is calculated using the sequence-dependent expected error and the read coverage at that site.

### Statistical analysis

All experiments were performed in multiple, independent experiments, as indicated in the figure legends. All statistics and tests are described fully in the text or figure legend.

## Code availability

Scripts for all analyses presented in this paper, including all data extraction, processing, and graphing steps are freely accessible at https://github.com/RouhanifardLab/PsiNanopore.git.

## Data availability

All raw and processed data used to generate figures and representative images presented in this paper are available at https://www.ebi.ac.uk/ena/browser/view/PRJEB56934.

## Acknowledgments

SHR acknowledges support from a Seed Networks Award from the Chan Zuckerberg Initiative CZF2019-002424 and NIH 5R01HG011087-02. MW acknowledges support from NIH R01HG10087 and Oxford Nanopore Technologies. YMH acknowledges support from NIH HG011120.

## Author Contributions

ST, MW and SHR conceived of the research. ST designed and performed the experiments. ST, MN, AM, CAM and NR analyzed the data with guidance from MW and SHR. HG designed and synthesized synthetic RNA controls with guidance from YMH. ST wrote the paper with guidance from YMH and SHR.

