## Supplementary figures for "Semi-quantitative detection of pseudouridine modifications and type I/II hypermodifications in human mRNAs using direct and long-read sequencing"

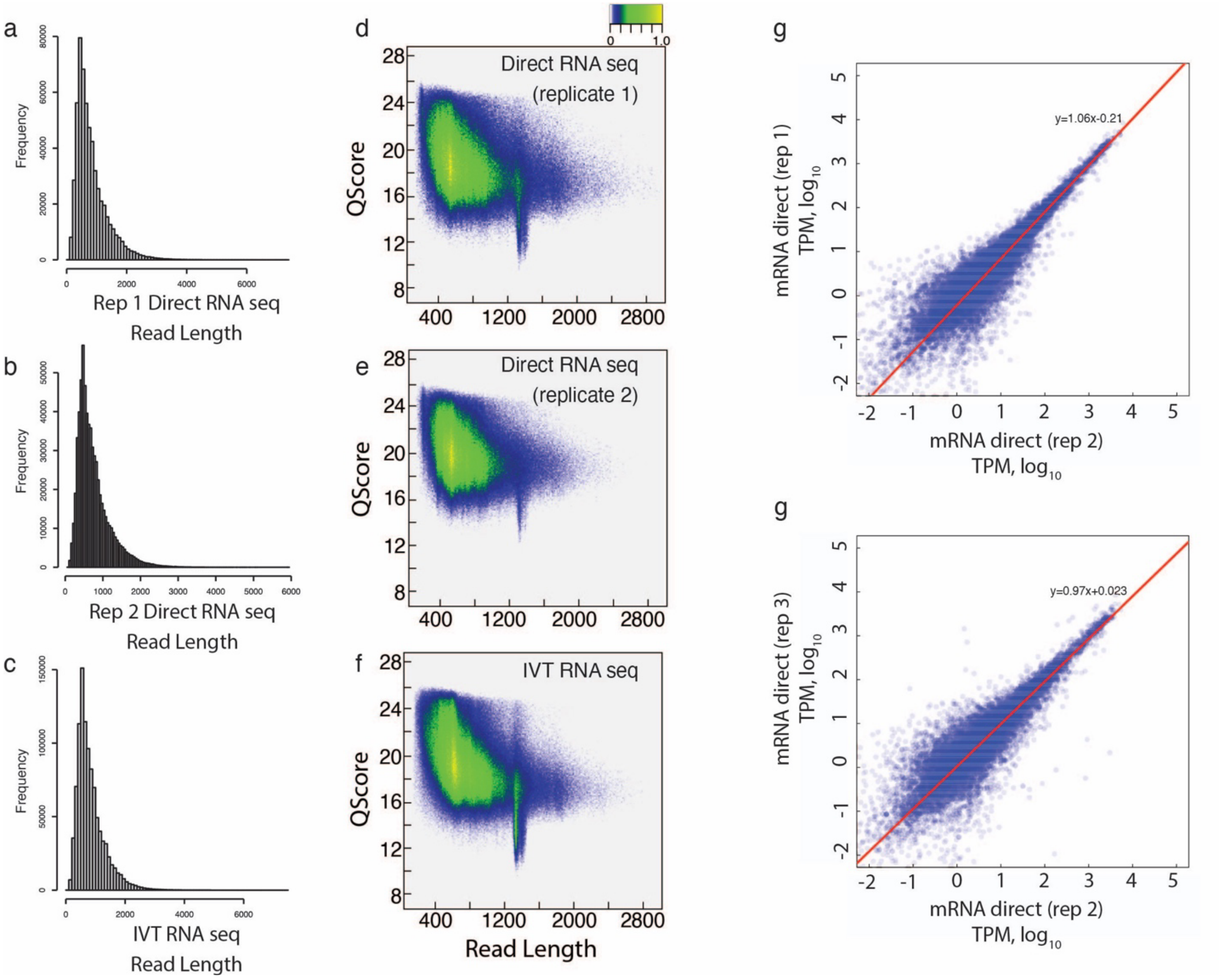


**Supplementary Figure 1: Analysis of replicates. a**, **b, c.** Read length distribution for $1^{st}$ and $2^{nd}$ replication of direct poly-A RNA sequencing and *in vitro* transcribed control (IVT) **d,e,f.** The read quality scores for the $1^{st}$ and $2^{nd}$ replicates of direct poly-A RNA sequencing and *in vitro* transcribed control (IVT).**g.** Correlation analysis of individual genes between replicates 1 and. 2 of direct RNA sequencing. **h.** Correlation analysis of individual genes between replicates 1 and 3.

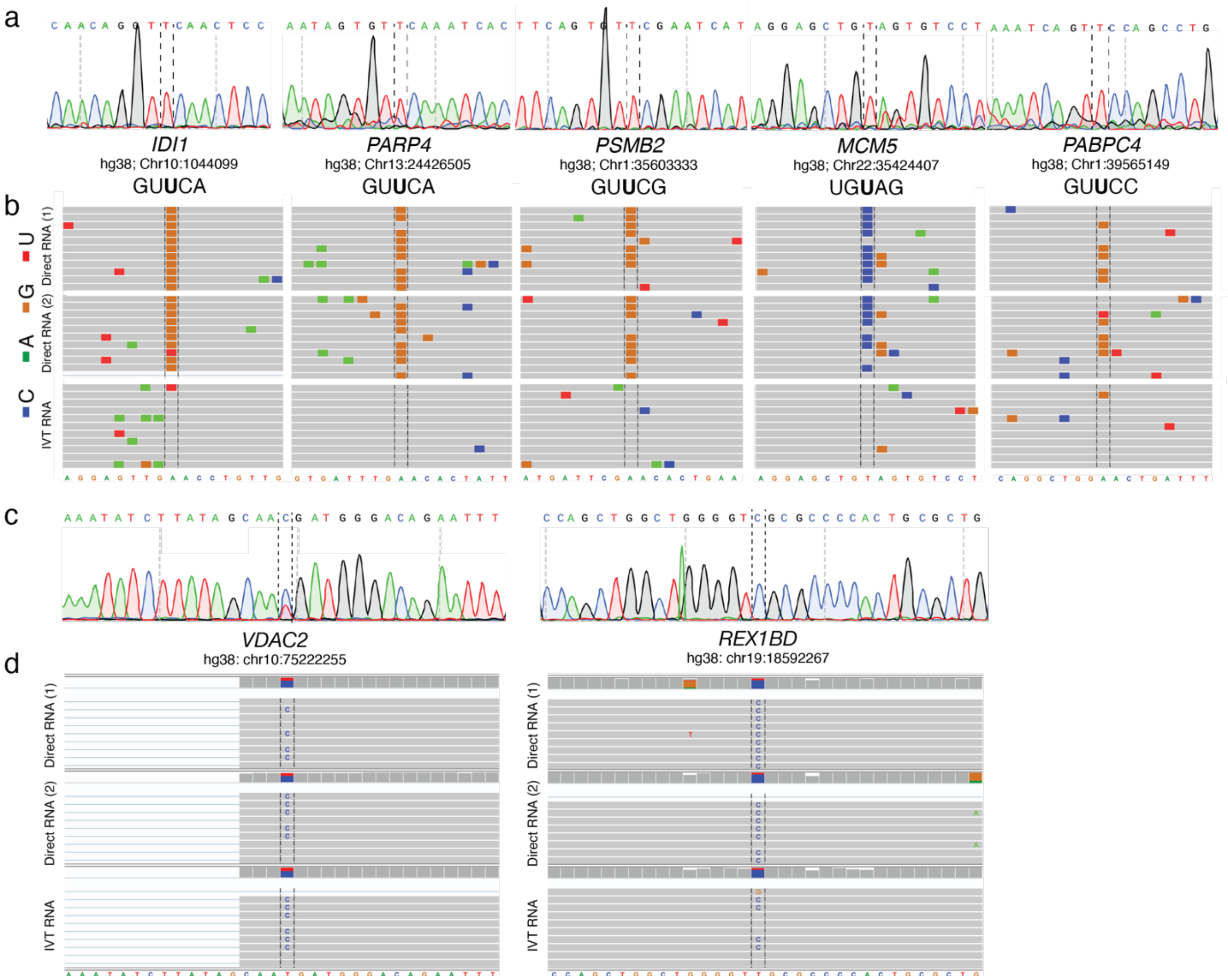


**Supplementary Figure 2: Sanger sequencing of HeLa gDNA for previously validated pseudouridylated positions and positions containing SNP. a,b.** Sanger sequencing results of HeLa gDNA for the positions that has been validated by previous detection methods(a) with the IGV snapshot of the corresponding region(b). c,d. Sanger sequencing result of HeLa gDNA and IGV snapshots of two positions *VDAC2* (chr10:75222255) and *REX1BD* (chr19:18592267) that contain and SNP in the middle (c) with the IGV snapshot of the corresponding region .

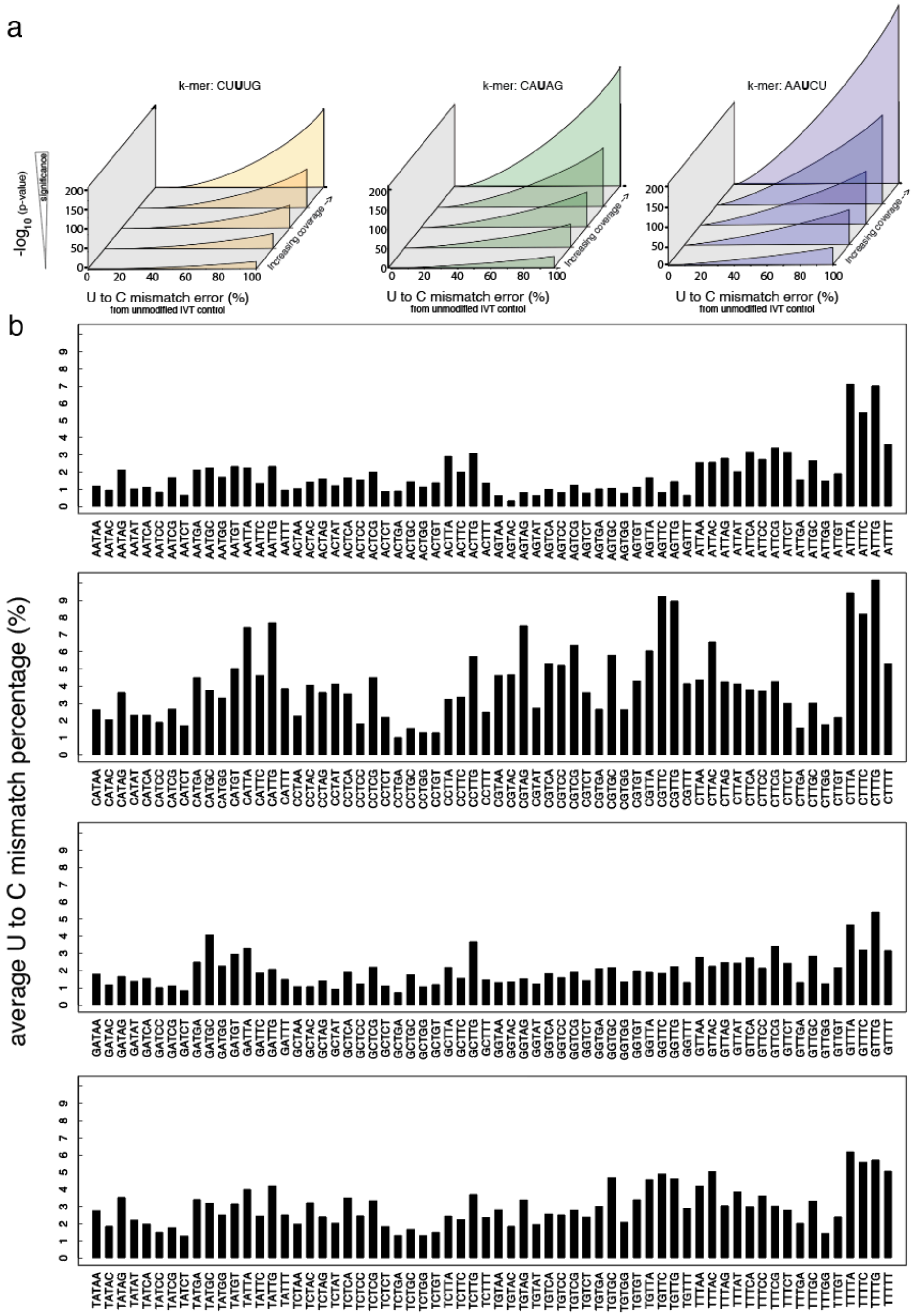


**Supplementary Figure 3: The average U to C mismatch percentages for kmers contain U in the middle. a.** Significance for different U-to-C mismatch percentages, and read coverages, for a high-error k-mer (CUUUG), medium-error k-mer (CAUAG) and a low-error k-mer (AAUCU).**b.** The average mismatch for each different 5mer that contains a U in the middle is shown.

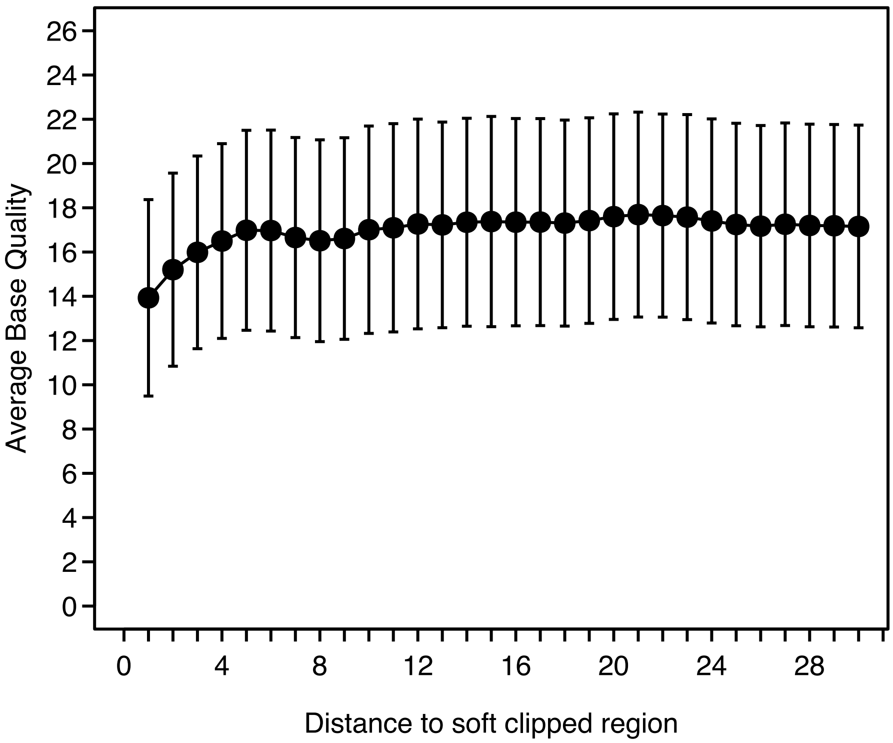


**Supplementary Figure 4: The average base quality versus the distance to soft clipped region.** The average base Quality vs the distance to soft clipped region for IVT control library.

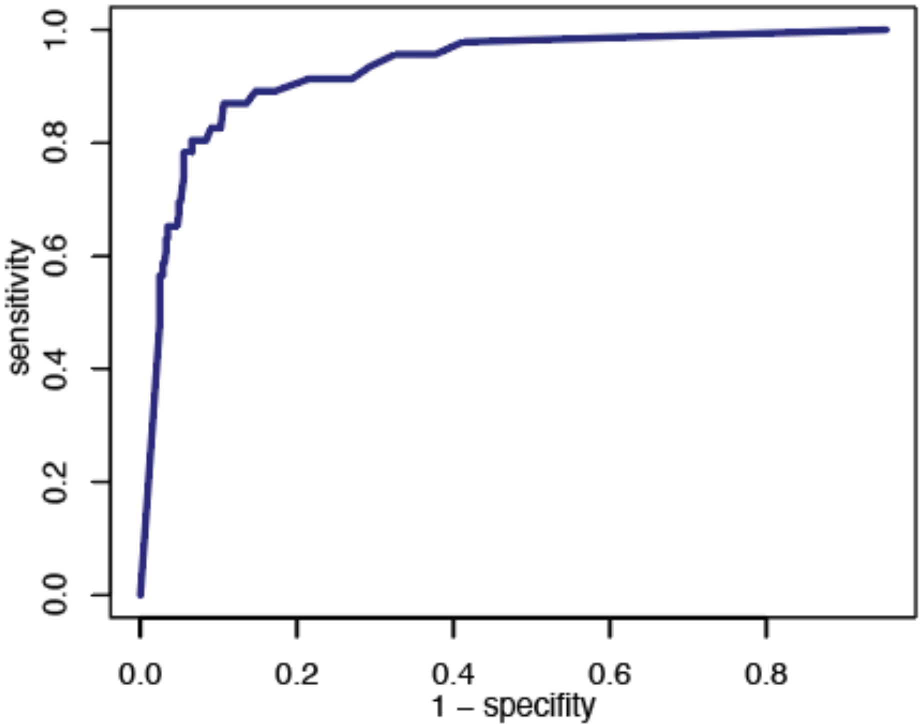


**Supplementary Figure 5. ROC curve for rRNA**
ROC curve comparing the validated pseudouridine targets detected in rRNA to the sites with p <0.001 using our method.

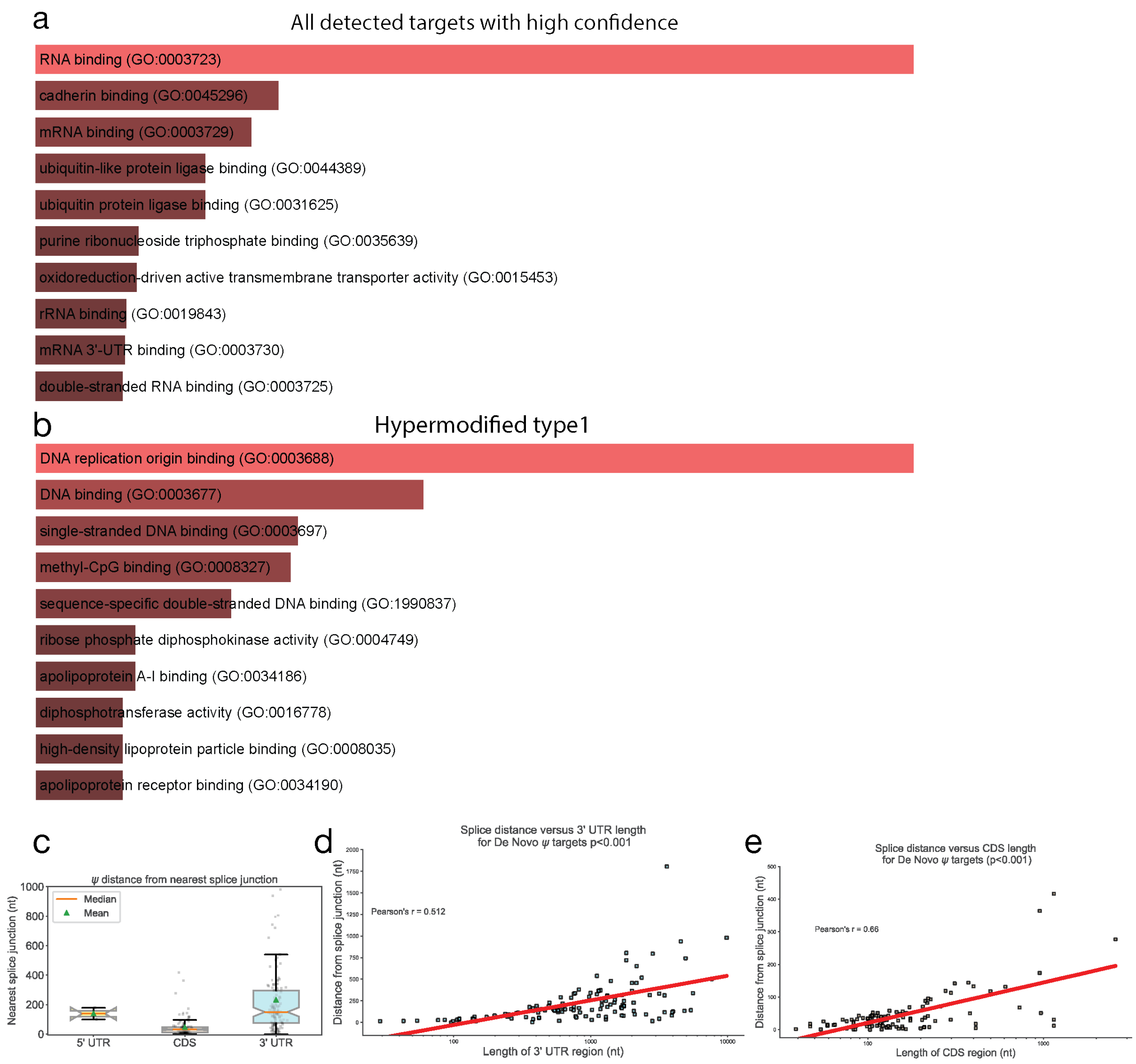


**Supplementary Figure 6. GO analysis and gene location analysis of de novo psi modifications**. a. The gene ontology (GO) analysis of the targets that are detected by nanopore method with high confidence (p-value < 0.001) b. The gene ontology (GO) analysis of the hypermodified type1 targets (mismatch differences of higher than 40% between direct and IVT) that are detected by nanopore method with high confidence and (p-value < 0.001) c. The distance from the nearest splice junction of the sites detected in the 5'UTR, 3'UTR, or CDS after reads were assigned to a dominant isoform using FLAIR [37] d. Correlation of the distance between the nearest splice site and targets located on the 3'UTR region versus the full length of that particular 3'UTR. e. Correlation of splice distance of targets located on a CDS region of their respective dominant isoform versus the full length of that particular CDS

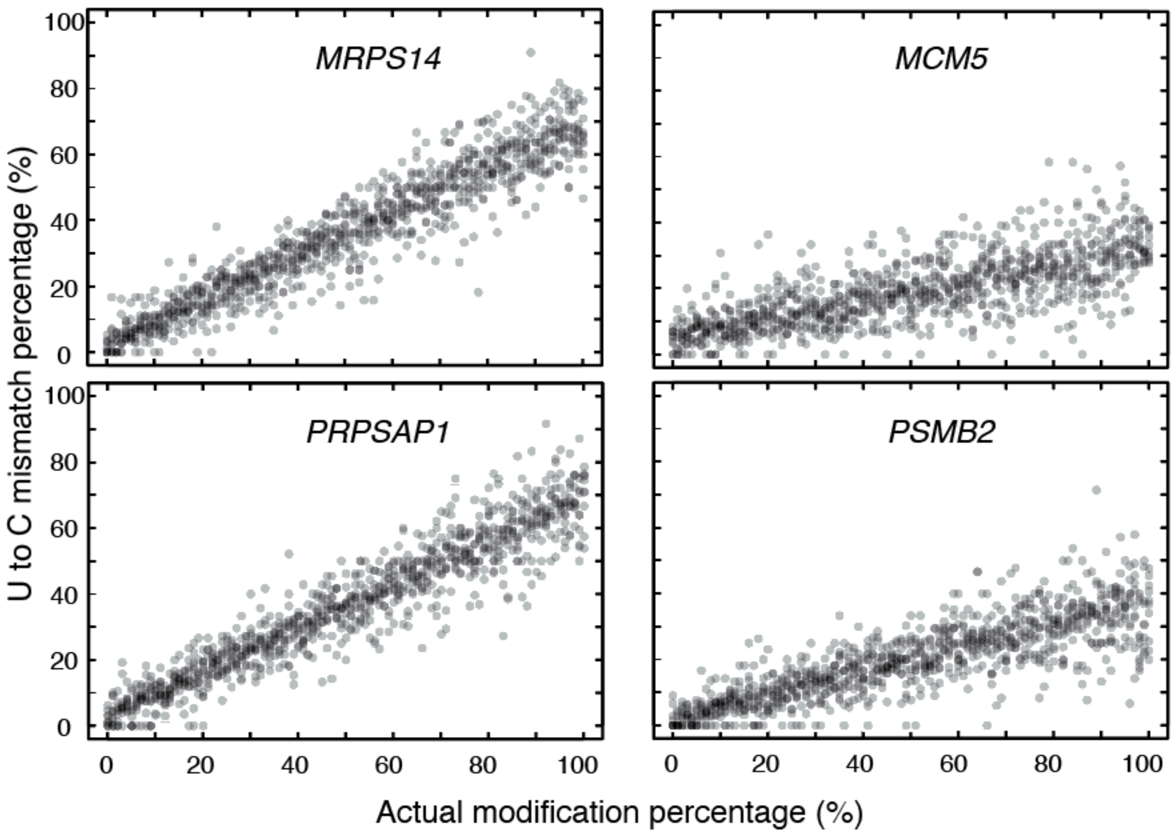


**Supplementary Figure 7. Synthetic transcripts experiment to see bias for lower percentages.** The Actual modification percentage of synthetic oligos versus observed U to C mismatch percentage.

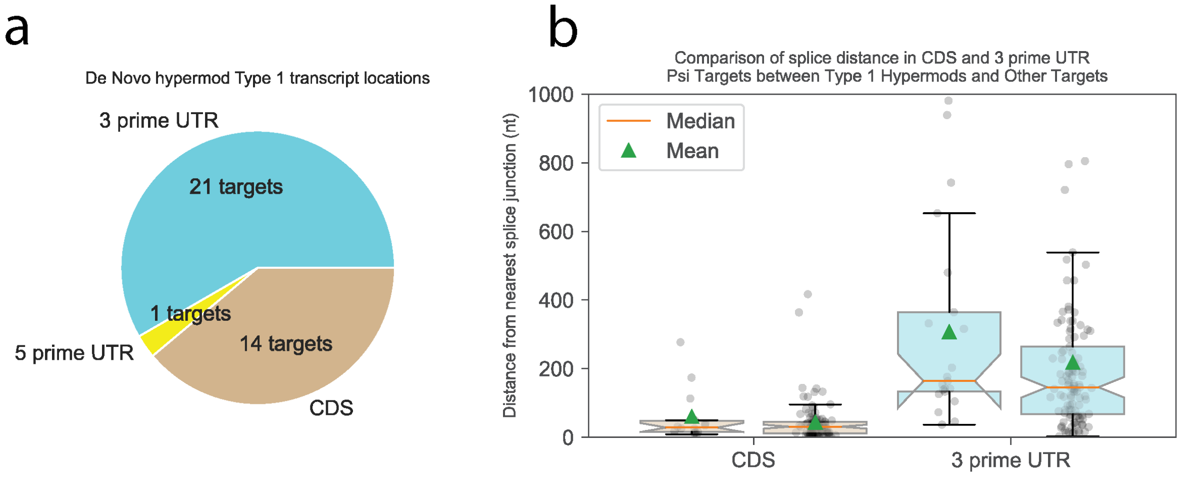


**Supplementary Figure 8. Hypermod type 1 location on gene and distance to splice junction. a.** The pie chart of transcriptome location of Type1 hypermodified targets. **b.** The box plot showing the distance from nearest splice junction for the targets detected in 3'UTR or CDS.
